# An investigation of carbapenemase-encoding genes in *Burkholderia cepacia* and *Aeromonas sobria* nosocomial infections among Iraqi patients

**DOI:** 10.1101/2024.11.28.625853

**Authors:** Mushtak T. S. Al-Ouqaili, Rawaa A. Hussein, Bushra A. Kanaan, Ahmed T.S. Al-Neda

## Abstract

*Burkholderia cepacia* and *Aeromonas sobria* are difficult to eradicate due to their innate resistance to a variety of medications, and cause various diseases. The aim of this study was to investigate the occurrence of carbapenemase genes and patterns of antibiotic resistance in isolates of *B. cepacia* and *A. sobria.* A cross-sectional study was conducted in the Ramadi Teaching Hospitals in the Al-Anbar Governorate in 2024. Various study samples, were used to collect the studied bacteria. The antibiotic resistance was detected by the VITEK®2 System. The presence of carbapenemase genes was confirmed via PCR technique. In this investigation, seventy-five (75) isolates of *A. sobria* and *B. cepacia* were assessed. Of these, *A. sobria* made up 16.6% (n = 20) while *B. cepacia* accounted for 45.8% (n = 55). The study isolates showed highest antimicrobial resistance to piperacillin, cefepime, ceftriaxone (100%), ceftazidime (97.3%), and lowest antimicrobial resistance to imipenem (36%). The result showed 55/57 *recA* gene positive for differentiated *B. cepacia complex* from other Burkholderia spp. The overall prevalence of carbapenemase genes was 92.8%% (52/56) with *bla_KPC_*accounting for 80.8% (42/52) and *bla_GES_* for 19.2% (10/52) of the total. The 42 *B. cepacia* isolates that tested positive for carbapenem resistance were constituted of 38 *bla_KPC_* (n = 38) and two *bla_GES_* (n = 2); in contrast, four *bla_KPC_*(n = 4) and eight *bla_GES_* (n = 8) were present in the *A. sobria* isolates that tested positive for carbapenems resistance. None of isolates studied tested positive for the *bla_IMP_*gene. The recent study concluded that *recA* gene identification was more sensitive and specific technique for detection *B. cepacia complex* isolates. There was a notable predominance of *bla_KPC_* and *bla_GES_*carbapenemase producers among the isolates under investigation. The *bla_IMP_*gene was not found in any of the research isolates.

## Introduction

*Burkholderia cepacia* has phenotypic characteristics in common with other members of the Burkholderiaceae family (previously known as *Pseudomonas cepacian*). These gram-negative bacteria are rod-shaped, motile, free-living, and range in size from 1.6 to 3.2 μm. It has been discovered that they have attachment-related pili in addition to multitrichous polar flagella. Soil, water, and plants, animals, and people in poor health can all contain *B. cepacia*. [1]. The nosocomial opportunistic bacterium *B. cepacia*, can seriously infect people with compromised immune systems. Because of their high degree of resistance to numerous treatments, their ability to tolerate a broad range of physical conditions, and their high transmissibility outside the hospital setting, *B. cepacia* strains are very persistent in the community, as well, of course, as within the hospital setting itself [2]. According to Wigfield et al. (2002), the primary cellular resistance mechanisms that are primarily involved in the overall process of antibiotic resistance development in *B. cepacia* species include altered medication targets, enzymatically changed antimicrobials, permeability of the outer membrane (OMP), and efflux pumps [3]. *B. cepacia* infections are difficult to treat because the isolates from these infections typically have high levels of inherently resistance to particular antibiotics or classes of drugs, and without requiring gene increase or mutation to several antimicrobials, which include ticarcillin, cephalosporins, phosphoric antibiotics, polymyxin, and aminoglycosides [4]. Furthermore, Nzula et al. (2002) found that the *B. cepacia* complex cells vary in their inherent antimicrobial resistance models, which is most likely related to the genomovar type [5].

*Aeromonas* species are gram-negative heterotrophic bacteria that are primarily found in warm areas and can contaminate water, seafood, meat, and vegetables, and which can lead to human illness [6]. *Aeromonas* infections range in frequency from 20 to 76 cases per 1,000,000 people. *A hydrophila, A. sobria,* and *A. caviae* are the most frequently isolated species [7]. These bacteria can cause a wide range of infections in humans, including of the gastrointestinal system, septicemia, acute respiratory tract infections, and soft tissue infections. Despite the fact that immunocompromised people are more likely to contract the disease, immunocompetent hosts can also contract it [8].

Because *Aeromonas* infections have at least three chromosomal β-lactamases, a major worry with such is their possible resistance to penicillin, ampicillin, carbenicillin, and cefazolin. Aeromonas spp. have been shown to produce Ambler class B, C, and D β-lactamases. The main β-lactamases that *Aeromonas* harbors are penicillinase, metallo-β-lactamases (MBL), and AmpC β-lactamases [9,10]. Worldwide, nosocomial infections are a problem for both developed and low-resource nations [11]. One of the main reasons for increased mortality and morbidity amongst people who have been hospitalized is infections contracted in healthcare facilities; they pose a serious risk to public health in general, as well as the sufferer in particular [12]. The aim of this study was to investigate the occurrence of carbapenemase genes and patterns of antibiotic resistance in isolates of *B. cepacia* and *A. sobria*.

## Materials and Methods

### Ethics Statement

The administrative human Ethical Approval Committee of the University of Anbar in Ramadi, Iraq, approved the study methods concerning the patients correspondingly to the official order numbered 91 in 22-5-2024 (valid from the date of application in 5-1-2024 according to the committee decision). During the study, the willingness of all participants to participate it was determined according to the Helsinki Declaration of 1979. Informed written consent was provided by all patients (or their parents) participating in the study.

### Collecting and Processing Specimens

From January to July 2024, patients who attended the Ramadi Teaching Hospitals in the Al-Anbar Governorate were randomly selected for collection of specimens, totaling 120 in all. The participants were aged between 15 and 60 years old, constituted of 50 (41.7%) females and 70 (58.3%) male participants. A questionnaire was used to gather data on the included patients. Many of the samples were cultivated on culture media such as MacConkey agar, Blood agar, and Mannitol agar (Oxoid, UK), as obtained from CSF, urine, ear swabs, burns, and wounds. These were subsequently incubated aerobically in a sterile environment at 37°C overnight [13].

### A. *cepacia and A. sobria* Isolate Identification

Each gram-negative isolate that was thought to have *B. cepacia* or *A. sobria* traits was identified via biochemical testing, oxidase and motility tests, microscopic and morphological characteristics, and streaking on culture media (MacFaddin 2000). Ultimately, any such diagnosis was confirmed via the VITEK-2 System (BioMérieux, France). Subsequent to this procedure, pure isolates were stocked in 20% glycerol incorporated in BHI (Oxoid, UK).

### Antibiotic Susceptibility

According to the criteria defined by the Clinical and Laboratory Standards Institute (CLSI) the antibiotic susceptibilities of *B. cepacia* and *A. sobria* isolates were tested using the VITEK-2 System (BioMérieux, Marcyl’Étoile, France) [11]. The antibiotics used included piperacillin, piperacillin/tazobactam, cefazolin, ceftazidime, ceftriaxone, cefepime, imipenem, meropenem, aztreonam, minocycline, amikacin, levofloxacin, ciprofloxacin, tigecycline, gentamicin, and trimethoprim-sulfamethoxazole.

### Methods for Phenotypic Identification of Metallo-β-Lactamase (MBL) production

A total of 56 strains resistant to at least one carbapenem were investigated. In this experiment, a disc containing meropenem (10 μg) and meropenem-EDTA (MBL inhibitor solution (0.5 M EDTA) (10/750 μg)) was applied to the lawn culture of the test strain’s 0.5 McFarland inoculum. After incubating for 16–18 hours at 35°C, the zones of inhibition of the two discs showed a difference in diameter of ≥ 7 mm, which suggested MBL formation [14].

### DNA Extraction

After cultivating pure single colonies of *B. cepacia* and *A. sobria* isolates on brain heart infusion broth overnight under strictly sterile conditions, the entire DNA of 75 *B. cepacia* and 20 *A. sobria* isolates was extracted using specific DNA extraction kits (Sacace, Italy), in accordance with the directions and guidelines of the manufacture company [15,16]. All nucleic acids were collected and frozen at below -20°C using a deep freezer. The PCR technique was then utilized to search for, and identify, all the genes listed in Table 1, 2.

**Table 1.**
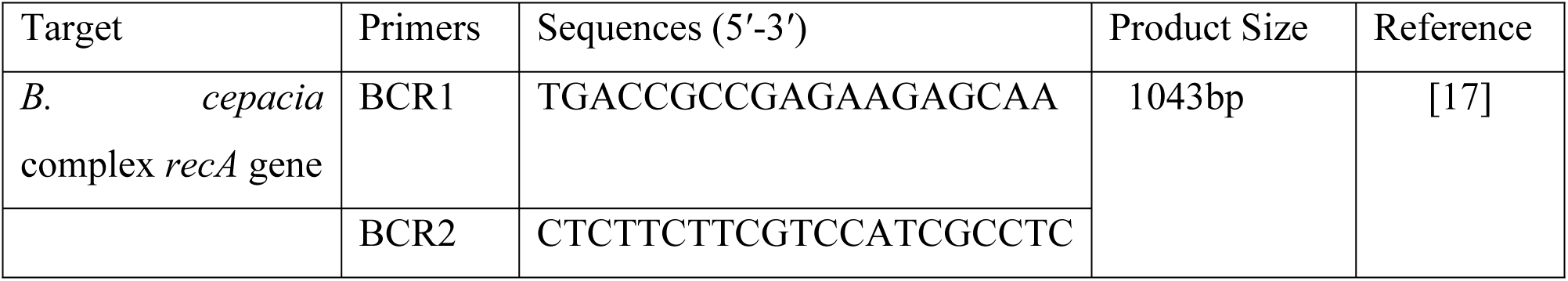
Primers used in this study for the detection of the *recA* gene.

### PCR techniques for *recA* gene detection in *B. cepacia* isolates

Using the *recA* gene of the Bcc complex in clinical samples and BCR1 and BCR2 primers, a standard PCR technique was used to directly molecularly identify the Bcc complex in all 75 extracted DNA samples (Table 1). A 25 μl reaction system comprising 12.5 μl of GoTaq® Green Master Mix (2X) (Promega, USA), 1 μl of each forward and reverse primer, 7.5 μl of nuclease-free water, and 3 μl of extracted DNA template was used for the PCR experiments. For the *recA* gene, the PCR reaction protocol was set up as follows: 30 cycles of 45 seconds at 94 ^O^ C, 45 seconds at 56 ^O^ C, and 90 seconds at 72 ^O^ C were followed by one cycle of 94 °C for two minutes. At 72°C, the last extension step was set for a single 7-minute cycle. Using a molecular size measurement of one hundred base pairs (from Bioneer, Korea), the PCR products were seen on a 1.5% agarose gel stained with red safe nucleic acid staining (Intron, Korea) [17].

### Molecular methods for identification of carbapenemase genes

12.5 μl of GoTaq® Green Master Mix (2X), 1 μl of forward and reverse primers (Macrogen, Korea), 3 μl of bacterial DNA, 0.5 μl of MgCl_2_, and 7 μl of double distilled and deionized water made up a total volume of 25 μl. The following conditions for PCR cycling were employed: 30 cycles of one minute each at 95°C, one minute at 50 or 55°C, one minute at 72°C, and ten minutes at 72°C (as indicated by Table 2). The PCR product was analyzed using a 100 bp molecular size marker gel and detected using Red Safe Nucleic Acid Staining [16] (Intron, Korea).

**Table 2.**
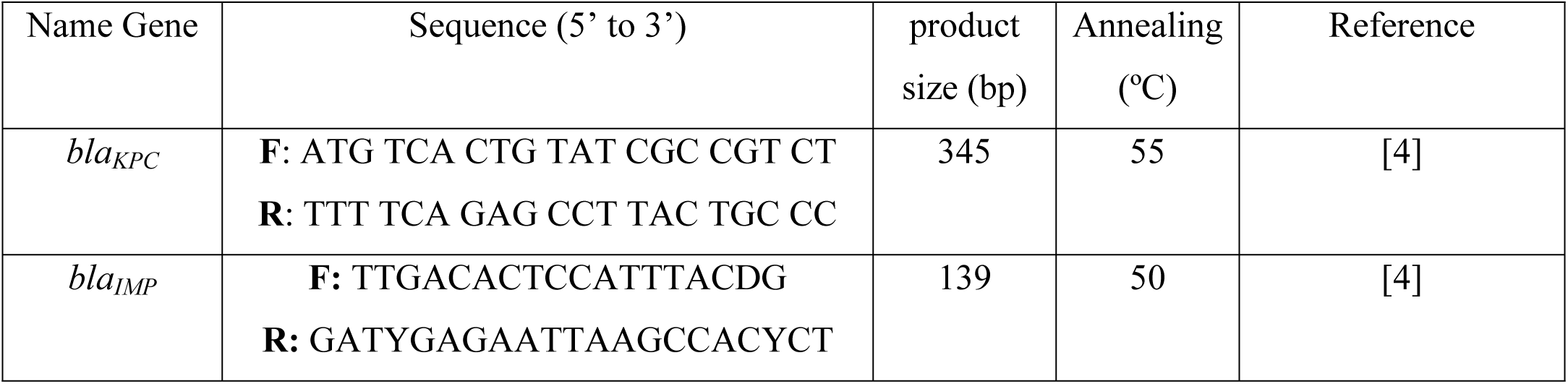
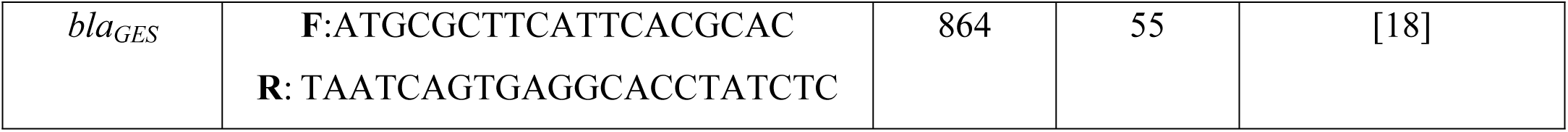
The specific primers are used in this investigation.

### Statistical Analysis

The results of the current work were analyzed via the chi-squared statistical method, as implemented in GraphPad version 10.2.0. Analysis of variance indicated the comparisons were statistically significant when P < 0.05.

## Results

### Features and Demographics of the Isolated Bacteria

In this investigation, seventy-five (75) isolates of *A. sobria* and *B. cepacia* were assessed. Of these, *A. sobria* made up 16.6% (n = 20) while *B. cepacia* accounted for 45.8% (n = 55). A majority of the study isolates (45.3%) were obtained from wound swabs, followed by urine (33.3%), diabetic foot ulcers (9.3%), and other sources (Table 3). The six (6) different sample types were used to obtain the study microorganisms. In terms of the gender distribution of the individuals from whom the isolates were collected, 44 (58.7%) were male and 31 (41.3%) female (see Table 3).The average age of the study participants, who ranged in age from 10 to 70, was 40.13 ± 22.5 years. Patients over 60 years old accounted for 34.6% of the isolates, followed by those between the ages of 21 and 30 (29.3%). The patients from whom the fewest specimens taken (4%) were those between the ages of 11 and 20.

**Table 3.** The distribution of study bacteria in relation to sample type, age, and gender of the subjects.

| Demographic Characteristics | Bacterial Isolates |  |  |  | Total (n, %) |
| --- | --- | --- | --- | --- | --- |
|  | *B. cepacia* |  | *A. sobria* |  |  |
|  | n. | % | n. | % |  |
| Gender |  |  |  |  |  |
| Male | 38 | 50.7% | 6 | 8% | 44 (58.7%) |
| Female | 17 | 22.6% | 14 | 18.7% | 31 (41.3%) |
| Age (years) |  |  |  |  |  |
| ≥10 | 5 | 6.7% | 1 | 1.3 % | 6 (8%) |
| 11–20 | 1 | 5.3 % | 2 | 2.6 % | 3 (4 %) |
| 21–30 | 14 | 18.6 % | 8 | 10.7% | 22 (29.3%) |
| 31–40 | 9 | 12 % | 1 | 1.3 % | 10 (13.3%) |
| 41–50 | 1 | 1.3 % | 3 | 4 % | 4 (5.3 %) |
| 51–60 | 3 | 4 % | 1 | 1.3 % | 4 (5.3 %) |
| > 60 | 22 | 29.3 % | 4 | 5.3 % | 26 (34.6%) |
| Type of sample |  |  |  |  |  |
| Wound swab | 26 | 34.6% | 8 | 10.6% | 34 (45.3%) |
| Urine | 18 | 24 % | 7 | 9.3 % | 25 (33.3%) |
| Ear swab | 1 | 1.3 % | 1 | 1.3 % | 2 (2.6 %) |
| Cerebrospinal fluid | 2 | 2.6 % | 0 | 0 % | 2 (2.6 %) |
| Diabetic foot ulcer | 5 | 6.7% | 2 | 2.7 % | 7 (9.3%) |
| Blood | 3 | 4 % | 2 | 2.7 % | 5 (6.7%) |

**P-value = 0.422 non-significant**

### Antimicrobial susceptibility

The isolates showed resistance to cefepime, piperacillin, and ceftriaxone (100%), ceftazidime (97.3%), and imipenem (36%). According to the VITEK-2 System, a total of 56 (74.6%) of the *B. cepacia* and *A. sobria* isolates were shown to exhibit carbapenem resistance. Of the 56 isolates, *B. cepacia* accounted for 75% (n = 42) and *A. sobria* for 25% (n = 14). Of the 56 isolates that tested positive for carbapenem resistance, 50.9% were resistant to meropenem and 36% to imipenem. Furthermore, of the 56 isolates resistant to carbapenem, 38 (67.8%) were collected from male patients, and 28 (50%) from female. The range of carbapenem MICs in this investigation was 0.2 μg/ml to 64 μg/ml. The results for meropenem MIC (which indicated that 36 isolates were resistant to carbapenem whilst 18 isolates were susceptible) appeared to be a more accurate indicator of the presence of the carbapenemase gene than the results for imipenem (which showed that 29 isolates with the carbapenemase gene were sensitive to carbapenem). Additionally, all seven isolates that harbored the carbapenemase gene were resistant to carbapenem. Profiles of drugs resistance are illustrated in Figure 1.

**Figure 1.**
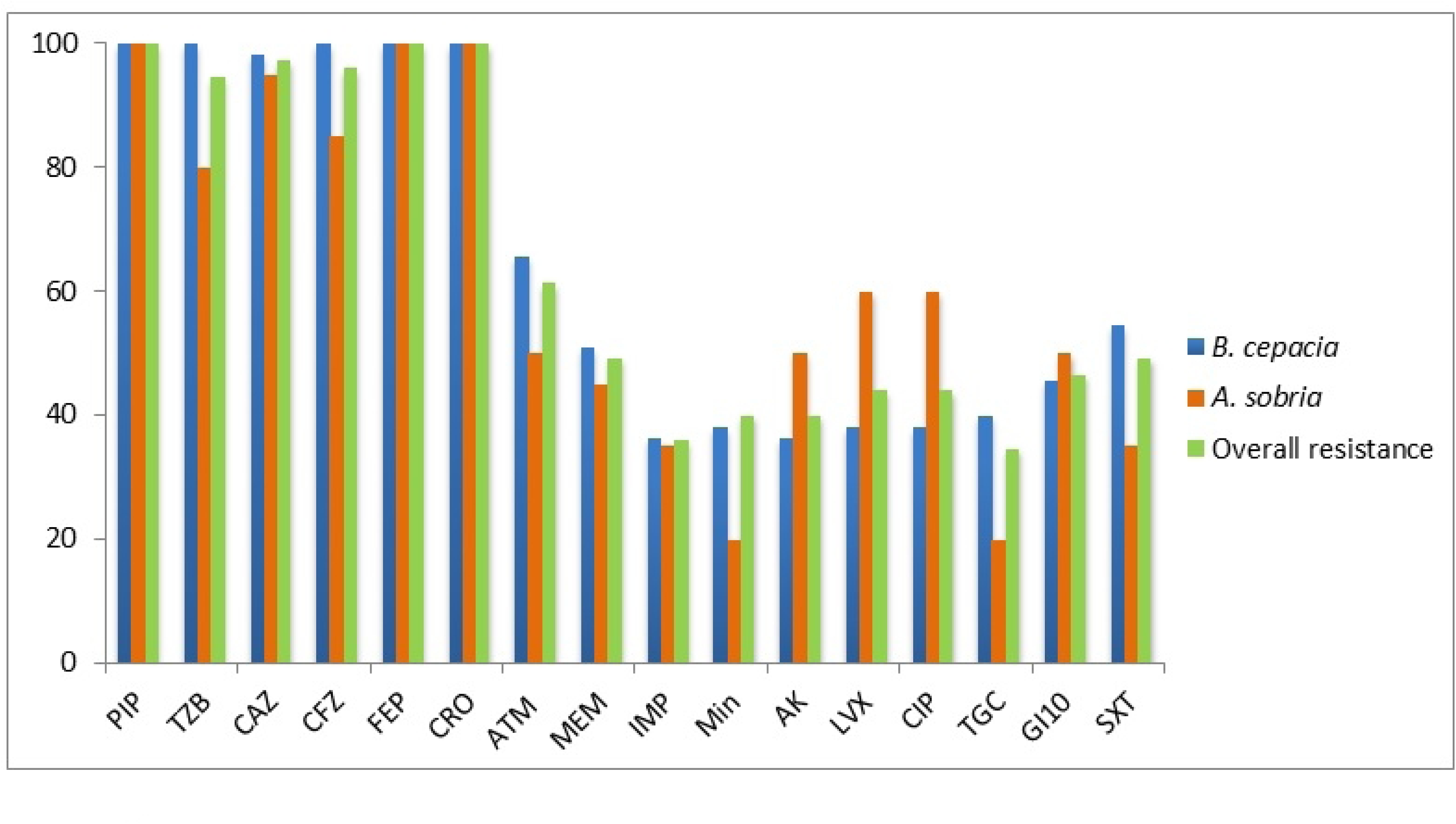
Patterns of drug resistance among the bacteria considered in the investigation. Depending on MICs, the antimicrobial susceptibility surveillance was detected for piperacillin (PIP), piperacillin/tazobactam (TZB), ceftazidime (CAZ), cefazolin (CFZ) cefepime (FEP), ceftriaxone (CRO), meropenem (MEM), imipenem (IMP), amikacin (AK), levofloxacin (LVX), ciprofloxacin (CIP), tigecycline (TGC), gentamicin (GI10), and trimethoprim-sulfamethoxazole (SXT).

### *recA* gene detection in *B. cepacia* isolates

The *recA* gene was utilized to identify the bacteria *B. cepacia*, and the study found that 55 out of 57 samples had the gene, which was identified using the VITEK 2 System device, as shown in figure 2. Four isolates were found to be *B. cepacia* complex by VITEK 2 System . Two isolates tested negative for *recA* gene and were found to be *B. cepacia* by VITEK 2 System .

**Figure 2.**
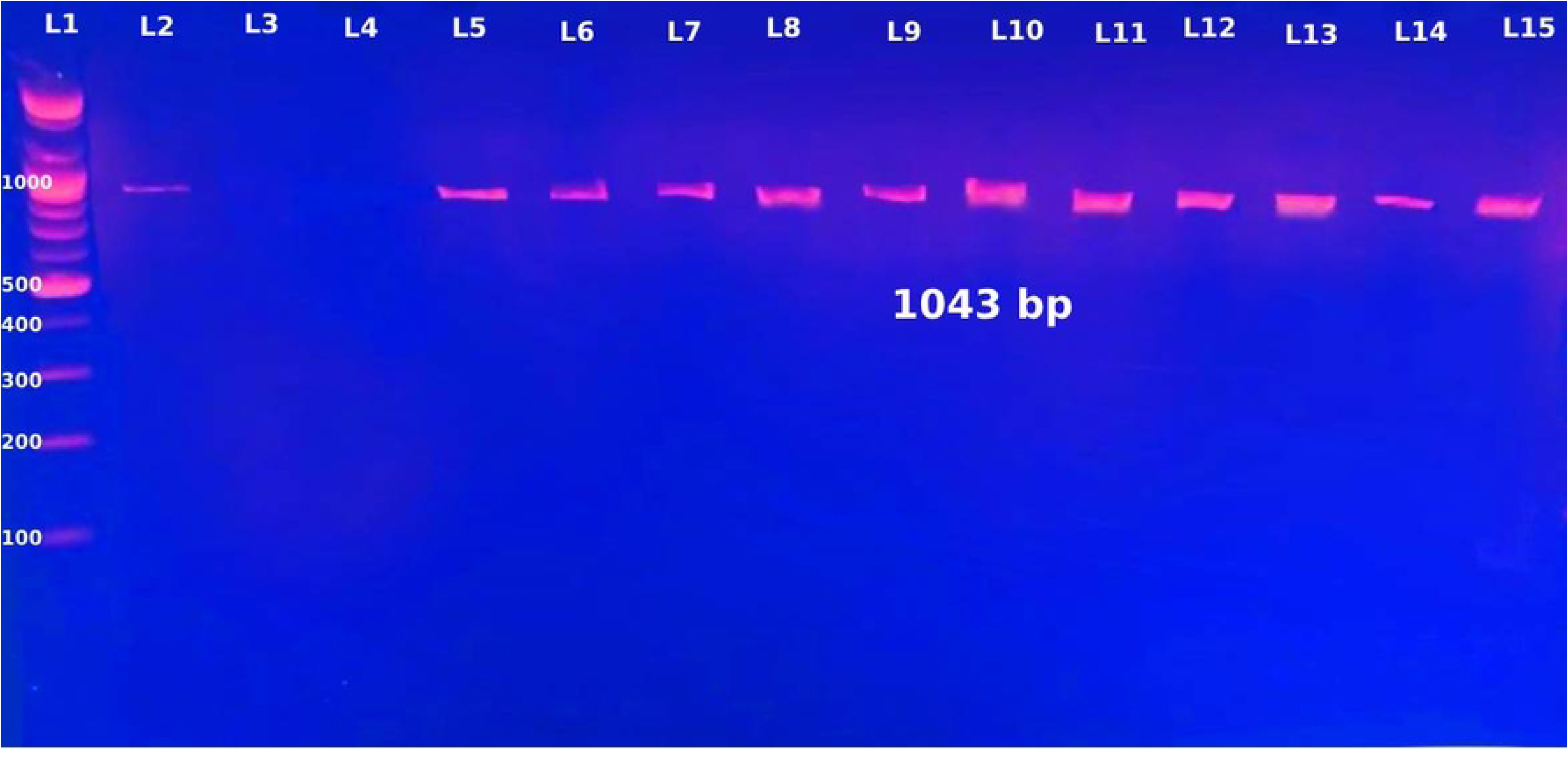
The recA gene’s gel electrophoresis (1043 base pairs). The lanes (2,4,5,6,7,8,9) were used to load the DNA samples. In lane 1, the molecular weight marker was the 100 base pair DNA ladder.

### Phenotypic Validation of the Synthesis of Carbapenemase and the Occurrence of Gene Encoding for Carbapenemase in Isolates of *B. cepacia* and *A. sobria*

In terms of the phenotypic validation of carbapenem resistance, the double disk synergy test yielded positive results for 87.5% (49/56) of the isolates that were resistant to meropenem and imipenem via double disk synergy (Table 4 and Figure 3). The double disk synergy test and PCR were used to determine the meropenem and imipenem resistance characteristics as mediated by the *bla_KPC_* and *bla_GES_* genes. No evidence for the *bla_IMP_* gene was found. Tables 5 and 6 report the distribution of the various carbapenemase genes. PCR was used in this investigation to identify the *bla_KPC_*-, *bla_IMP_*-, and *bla_GES_*-like genes among the 56 isolates resistant to carbapenem. Using PCR, the overall prevalence of carbapenemase genes was found to be 92.8%% (52/56) with *bla_KPC_*, accounting for 80.8% (42/52), and *bla_GES_*, for 19.2% (10/52) of the total. Of the 42 *B. cepacia* isolates that tested positive for carbapenem resistance, 38 *bla_KPC_*(n = 38) and two *bla_GES_* (n = 2) were found; in contrast, four *bla_KPC_* (n = 4) and eight *bla_GES_* (n = 8) were present in the *A. sobria* isolates that tested positive for carbapenems. There was no gene present in two isolates that were resistant to carbapenem. None of the isolates contained the *bla_IMP_* gene, as can be seen in Figures 4, 5. There was no gene present in the two isolates resistant to carbapenem. Additionally, no isolate was found in which carbapenemase genes were noted to coexist. 24 wound samples, 14 urine samples, and two CSF fluids tested positive for the total of 40 *bla_KPC_*- positive *B. cepacia*. *bla_GES_*, however, were found in the isolates gained from two diabetic foot ulcers. Four *bla_KPC_*-positive *A. sobria* were isolated from two samples from wounds and one sample each from urine and ears. *bla_GES_* were identified in urine (two), blood (three), and diabetic foot ulcers (three). The statistical analysis showed that resistance to imipenem, meropenem, cefepime, ceftriaxone, piperacillin (P-value = 0.000), piperacillin/tazobactam (P-value = 0.003), ceftazidime (P-value = 0.001), and cefazolin (P-value = 0.002) was significantly correlated with the presence of carbapenemase genes. The range of carbapenem MICs in this investigation was 0.2 μg/ml to 64 μg/ml.

**Figure 3.**
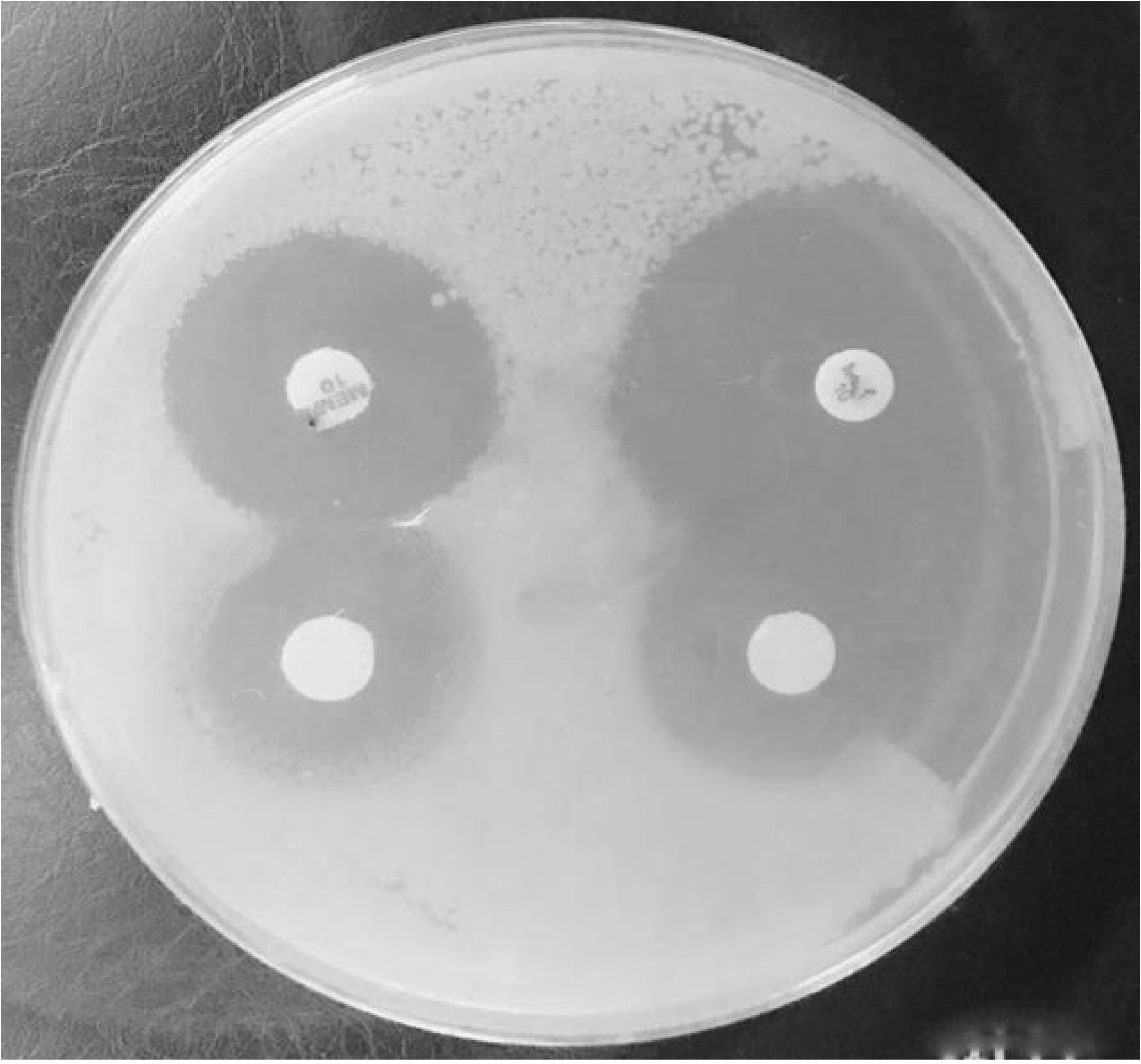
Phenotypic preliminary test for detection of carbapenemase production.

**Figure 4.**
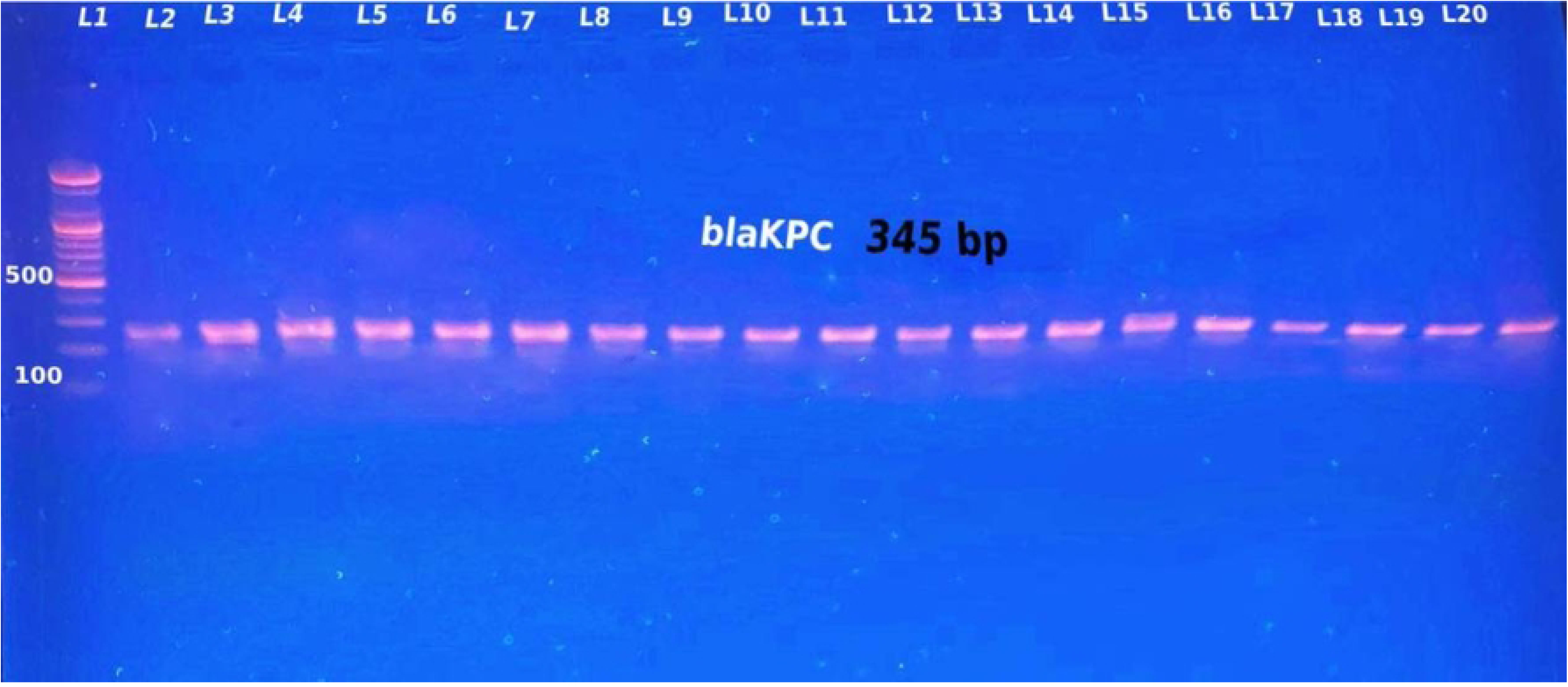
Positive results for the *bla_KPC_* gene were obtained via 1.5% agarose gel electrophoresis. The gene sizes in lanes 1–20 were found to be 345 bp. A DNA ladder in the first lane functioned as a 100 bp molecular weight marker

**Figure 5.**
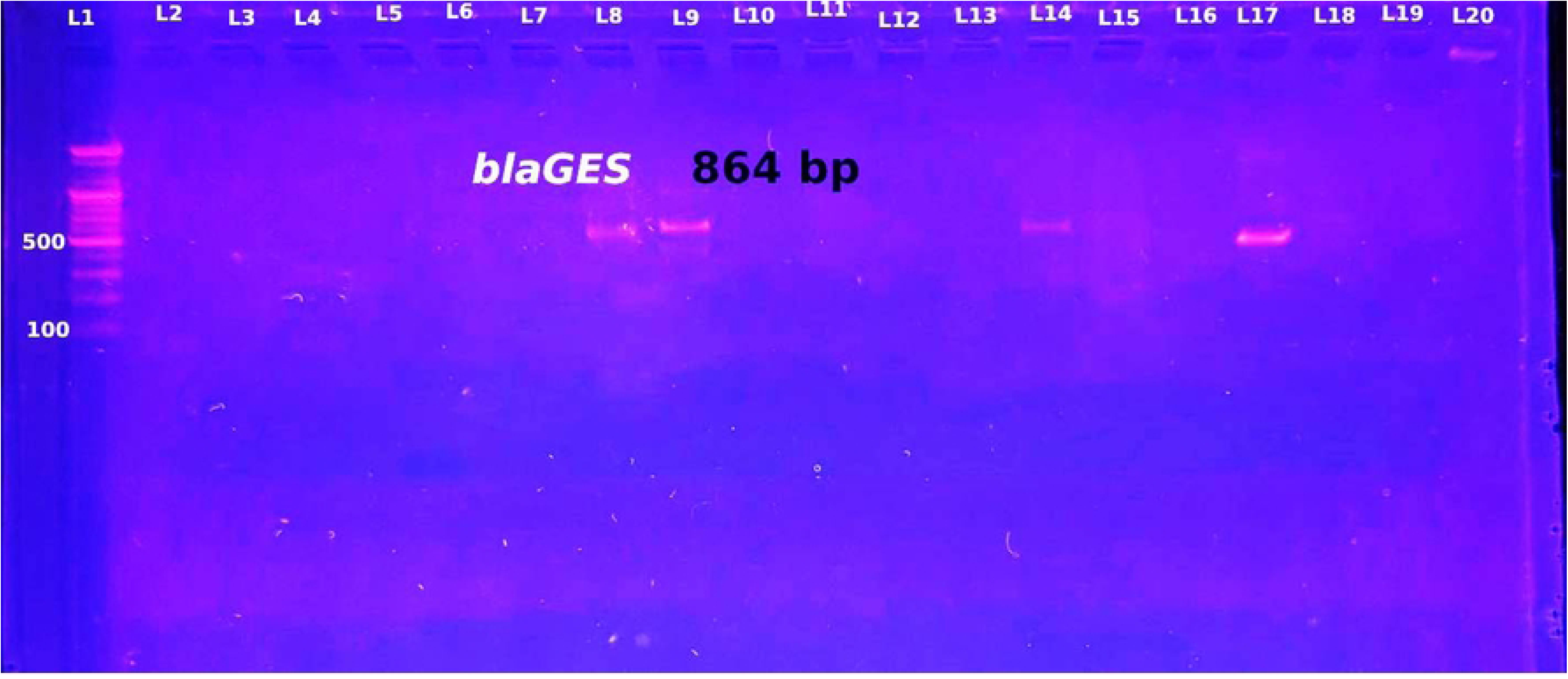
Positive results for the *bla_GES_* gene were obtained via 1.5% agarose gel electrophoresis. In lanes 8, 9, and 17, the gene sizes were 864 bp. A DNA ladder in lane 1 functioned as a 100 bp molecular weight marker.

**Table 4.** Findings regarding the phenotypic and genotypic methods for identifying carbapenem resistance.

| Isolates | Phenotype |  | Genotype |  |  |  |
| --- | --- | --- | --- | --- | --- | --- |
|  | Disc | Double disk synergy test | PCR | *bla<sub>KPC</sub>* | *bla<sub>IMP</sub>* | *bla<sub>GES</sub>* |
|  | n = 75 | n = 56 | n = 56 |  |  |  |
| *B. cepacia* | n = 42 (56%) | n = 39(69.6%) | 40 (71.4%) | 38 (67.9%) | - | 2 (3.6%) |
| *A. sobria* | n = 14 (18.6%) | n = 10 (17.9%) | 12 (21.4) | 4 (7.1%) | - | 8 (14.3%) |
| Total | n = 56 (74.6%) | n = 49 (87.5%) | 52 (92.8%) | 42 (75%) | - | 10 (17.9%) |

**Table 5.** Antibiogram and resistance genes of isolates of *B. cepacia* that were determined to be carbapenemase positive.

| Isolate No | Isolates | PCR | Carbapenems (MIC–µg/ml) |  |
| --- | --- | --- | --- | --- |
|  |  |  | MEM | IMP |
| 1 | *Burkholderia cepacia* | *bla<sub>KPC</sub>* | R (16) | R (16) |
| 3 | *Burkholderia cepacia* | *bla<sub>KPC</sub>* | R (32) | R (64) |
| 7 | *Burkholderia cepacia* | *bla<sub>KPC</sub>* | R (32) | R (32) |
| 8 | *Burkholderia cepacia* | *bla<sub>KPC</sub>* | R (16) | R (32) |
| 9 | *Burkholderia cepacia* | *bla<sub>KPC</sub>* | R (16) | R (16) |
| 10 | *Burkholderia cepacia* | *bla<sub>KPC</sub>* | R (64) | R (16) |
| 11 | *Burkholderia cepacia* | *bla<sub>KPC</sub>* | R (32) | S (0.5) |
| 12 | *Burkholderia cepacia* | *bla<sub>KPC</sub>* | R (32) | S (0.5) |
| 13 | *Burkholderia cepacia* | *bla<sub>KPC</sub>* | R (32) | S (0.5) |
| 14 | *Burkholderia cepacia* | *bla<sub>KPC</sub>* | R (16) | S (0.5) |
| 20 | *Burkholderia cepacia complex* | *bla<sub>KPC</sub>* | R (16) | S (4) |
| 21 | *Burkholderia cepacia complex* | *bla<sub>KPC</sub>* | R (32) | S (4) |
| 22 | *Burkholderia cepacia complex* | *bla<sub>KPC</sub>* | R (32) | S (0.2) |
| 23 | *Burkholderia cepacia complex* | *bla<sub>GES</sub>* | R (16) | S (0.2) |
| 24 | *Burkholderia cepacia* | *bla<sub>KPC</sub>* | R (16) | S (0.2) |
| 25 | *Burkholderia cepacia* | *bla<sub>KPC</sub>* | R (64) | S (4) |
| 26 | *Burkholderia cepacia* | *bla*<sub>KPC</sub> | R (16) | S (0.2) |
| 27 | *Burkholderia cepacia* | *bla*<sub>KPC</sub> | R (16) | S (4) |
| 28 | *Burkholderia cepacia* | *bla*<sub>KPC</sub> | R (16) | S (4) |
| 29 | *Burkholderia cepacia* | *bla*<sub>KPC</sub> | R (32) | S (4) |
| 35 | *Burkholderia cepacia* | *bla*<sub>KPC</sub> | R (16) | S (0.2) |
| 36 | *Burkholderia cepacia* | *bla*<sub>KPC</sub> | R (32) | S (0.2) |
| 37 | *Burkholderia cepacia* | *bla*<sub>GES</sub> | R (64) | S (0.2) |
| 38 | *Burkholderia cepacia* | *bla*<sub>KPC</sub> | R (16) | S (0.2) |
| 39 | *Burkholderia cepacia* | *bla*<sub>KPC</sub> | R (16) | S (0.2) |
| 40 | *Burkholderia cepacia* | *bla*<sub>KPC</sub> | R (16) | S (0.2) |
| 41 | *Burkholderia cepacia* | *bla*<sub>KPC</sub> | R (32) | S (0.5) |
| 42 | *Burkholderia cepacia* | *bla*<sub>KPC</sub> | R (32) | S (0.5) |
| 43 | *Burkholderia cepacia* | *bla*<sub>KPC</sub> | S (0.5) | R (16) |
| 44 | *Burkholderia cepacia* | *bla*<sub>KPC</sub> | S (0.2) | R (16) |
| 45 | *Burkholderia cepacia* | *bla*<sub>KPC</sub> | S (4) | R (16) |
| 46 | *Burkholderia cepacia* | *bla*<sub>KPC</sub> | S (1) | R (16) |
| 47 | *Burkholderia cepacia* | *bla*<sub>KPC</sub> | S (0.5) | R (16) |
| 48 | *Burkholderia cepacia* | *bla*<sub>KPC</sub> | S (0.5) | R (16) |
| 49 | *Burkholderia cepacia* | *bla*<sub>KPC</sub> | S (0.5) | R (16) |
| 50 | *Burkholderia cepacia* | *bla*<sub>KPC</sub> | S (0.5) | R (32) |
| 51 | *Burkholderia cepacia* | *bla*<sub>KPC</sub> | S (0.2) | R (32) |
| 52 | *Burkholderia cepacia* | *bla*<sub>KPC</sub> | S (0.2) | R (32) |
| 53 | *Burkholderia cepacia* | *bla*<sub>KPC</sub> | S (4) | R (32) |
| 54 | *Burkholderia cepacia* | *bla*<sub>KPC</sub> | S (1) | R (32) |
| 55 | *Burkholderia cepacia* | *bla*<sub>KPC</sub> | S (0.2) | R (64) |
| 56 | *Burkholderia cepacia* | *bla*<sub>KPC</sub> | S (0.2) | R (64) |

**Table 6.** Antibiogram and resistance genes of carbapenemase producing isolates of *A. sobria*.

| Isolate No | Isolates | PCR | Carbapenems (MIC-µg/ml) |  |
| --- | --- | --- | --- | --- |
|  |  |  | MEM | IMP |
| 2 | *Aeromonas sobria* | *bla*<sub>GES</sub> | S (0.2) | R (32) |
| 4 | *Aeromonas sobria* | - | S (0.5) | R (32) |
| 5 | *Aeromonas sobria* | *bla*<sub>GES</sub> | S (0.5) | R (16) |
| 6 | *Aeromonas sobria* | *bla*<sub>GES</sub> | S (4) | R (16) |
| 15 | *Aeromonas sobria* | - | R (16) | R (16) |
| 16 | *Aeromonas sobria* | *bla*<sub>KPC</sub> | R (16) | R (32) |
| 17 | *Aeromonas sobria* | *bla*<sub>GES</sub> | S (0.5) | R (64) |
| 18 | *Aeromonas sobria* | *bla*<sub>GES</sub> | R (32) | S (0.5) |
| 19 | *Aeromonas sobria* | *bla*<sub>KPC</sub> | R (32) | S (0.5) |
| 30 | *Aeromonas sobria* | *bla*<sub>GES</sub> | R (32) | S (0.5) |
| 31 | *Aeromonas sobria* | *bla*<sub>KPC</sub> | R (32) | S (0.2) |
| 32 | *Aeromonas sobria* | *bla*<sub>GES</sub> | R (64) | S (0.2) |
| 33 | *Aeromonas sobria* | *bla*<sub>GES</sub> | R (16) | S (1) |
| 34 | *Aeromonas sobria* | *bla*<sub>KPC</sub> | R (16) | S (1) |

## Discussion

In this investigation, seventy-five (75) *A. sobria* and *B. cepacia* isolates were assessed. Of these, *A. sobria* made up 16.6% (n = 20) while *B. cepacia* accounted for 45.8% (n = 55). In terms of the gender distribution of the individuals from whom the isolates were collected, 44 (58.7%) were male and 31 (41.3%) female. Out of the 44, *B. cepacia* detected in 50.7% of male patients while *A. sobria* was detected in 8%. With regard to the 31 female patients, *B. cepacia* was detected in 22.6% and *A. sobria* in 18.7%. The pattern of cases by gender was in good agreement with the literature, where it was found that isolates of *B. cepacia* were present in 46.70% of females and 53. 30% of males, respectively [18].

Pineda-Reyes and associates found that the distribution of *A. sobria* was 49.3% in males and 50.7% among females [19]. A majority of the study isolates (45.3%) were obtained from wound swabs, followed by urine (33.3%), diabetic foot ulcers (9.3%), and other sources. By contrast, 26 (34.6%) of the *B. cepacia* isolates were isolated from wound swabs, 18 (24%) from urine, five (6.7%) from diabetic foot ulcers, three (4%) from blood, two (2.6%) from cerebrospinal fluid, and one (1.3%) from ear swabs, according to the type of specimen. This result was consistent with Dizbay et al. (2009), who observed that *B. cepacia* is linked to a variety of human infections [20]. Eight (10.6%) of the *A. sobri* isolates were obtained from wound swabs, seven (9.3%) from urine, two (2.7%) from diabetic foot ulcers, two (2.7%) from blood, and one (1.3%) from ear swabs. Regardless of the type of specimen, no *A. sobria* was found in any of the cerebrospinal fluid specimens. This is consistent with Pineda-Reyes et al. (2024), who observed that *A. sobri* is linked to a variety of human infections [19,21].

Options for therapy with *B. cepacia* remain limited. Application of restricted medications such as Clavulanic acid, ceftazidime, meropenem, minocycline, levofloxacin, chloramphenicol, and trimethoprim-sulfamethoxazole is advised by the CLSI guide [11]. While resistances to ceftazidime and trimethoprim/sulfamethoxazole were 98.18% (54/55) and 54.54 (30/55), respectively, susceptibility to meropenem and minocycline remained more prevalent, with 40% (22/55) and 38.18 % (26/55) of isolates being resistant, respectively. The present study was in line with Al-Muhanna and associates, who found that isolates of *B. cepacia* exhibited a high degree of resistance to almost all of the β-lactam antibiotic classes they considered [21]. These included: 100% for ceftriaxone, cefoxitin, and cefepime; 87.5% for ticarcillin with Clavulanic acid, piperacillin, ceftazidime, tobramycin, ciprofloxacin, and levofloxacin; 62.5% for aztreonam and amikacin; and 37.5% for meropenem, imipenem, and gentamicin. The isolates of *A. sobri* in this investigation demonstrated little resistances to minocycline and tigecycline; 20% and 35%, respectively, to meropenem and imipenem; and high resistances to piperacillin, cefepime, and ceftriaxone (100%), and to ceftazidime (95%). The present study was in line with Rhee et al., who observed high antibiotic resistance (15.5%) for CRO and TZP, but only 9.8% and 3.0%, respectively, to amikacin and carbapenem [22].

In this work, *B. cepacia* and *A. sobria* isolated from clinical specimens in the Al-Anbar Governorate, Iraq, were evaluated for the prevalence of carbapenem resistance, AMR patterns, and carbapenemase gene distribution. This study is amongst the few, rare carbapenem resistance studies to consider this country. Resistances to meropenem and imipenem were 40% and 36.36% in *B. cepacian*, which appear to be higher than was found in Iraq by Abbas in 2016 [17] and Tuwaij, et al. [23]. For *A. sobria*, the frequency of meropenem and imipenem resistance was 35%, which was also greater than observed in ref. [24].

The first step in classifying an isolate within the B. cepacia complex is to amplify *recA* using primers BCR1 and BCR2. The recA gene was utilized to identify the bacteria *B. cepacia* complex and the study found that 55 out of 57 samples had the gene, which was identified using the VITEK 2 System device. Two isolates were found to be B. cepacia by VITEK 2 System and tested negative for recA gene. Brisse and colleagues discovered that, despite the fact that *B. gladioli* is not listed in the VITEK2 database (and so should have a higher probability of misidentification), only one isolate of the bacteria was recognized by the VITEK 2 apparatus as “*B. cepacia* or *B. pseudomalleii*.” However, as other isolates of these species or other non-fermenters like Achromobacter or Alcaligenes may be mistakenly recognized as *B. cepacia*, it is always an excellent choice to use a molecular approach to validate the *B. cepacia* identification [25]. The *recA* gene has been widely used in bacterial systematics and has proven to be highly helpful in identifying species of the *B. cepacia* complex. By using phylogenetic analysis of sequence variance within the gene, it is possible to distinguish between the nine present species of the *B. cepacia* complex. BCR1 and BCR2, the original recA-based PCR primers, are exclusive to the B. cepacia complex members and do not amplify this gene in other B. cepacia species. This limits the technique’s ability to classify other Burkholderia species in various natural habitats, even though it can be a useful method of confirming an isolate’s position within the complex. The gene technique (PCR) results are highly accurate, and we can rely on them [25].

Carbapenemases are enzymes that have various different hydrolytic profiles; all are categorized as β-lactamases. All the β-lactam antibiotics, including the penicillins, cephalosporins, monobactams, and the carbapenems, are acted upon by these enzymes. Due to carbapenemase activity, a majority of β-lactam drugs can become inactive against the severe infections that result from bacteria that produce such β-lactamases. Enzyme superfamilies’ comprise carbapenemases from the A, B, and D classes of β-lactamases. Zinc is present in the active region of class B enzymes, which are metallo-β-lactamases; class A and D enzymes, on the other hand, hydrolyze serine. The Guiana-Extended-Spectrum and *Klebsiella pneumoniae* carbapenemase families are amongst the Class A carbapenemases. The KPC carbapenemases, which are mainly found on *Klebsiella pneumoniae* plasmids, are the most well-known members of this category. The OXA-type β-lactamases are classified as class D carbapenemases, and are mostly generated by *Acinetobacter baumannii* [26]. Five further subgroups include IMP, VIM, SPM, GIM, and SIM, and mainly belong to *Pseudomonas aeruginosa*. However, an increasing number of cases of these kinds of β-lactamase in the Enterobacteriaceae have been reported globally. The characteristics, epidemiology, and detection of carbapenemases found in pathogenic bacteria have been updated in the present article [26,27]. Furthermore, the phenotypic confirmatory tests conducted in this study revealed that 87.5% (49/56) of the isolates had carbapenem resistance, namely via the double synergy test. Contrary to the current study, which concentrated on *B. cepacia* and *A. sobria* Codjoe et al. in Ghana revealed a significantly lower incidence of 18.9% via the phenotypic method, which may be because the majority of their isolates were Acinetobacter spp. and Pseudomonas spp [28,29]. Panduragan and associates, 2015 study in India indicated that 62% of isolates developed carbapenemases when both the phenotypic and genotypic approaches were used in gram-negative bacteria [30]. This finding is consistent with the current study, which identified similar proportions (87.5% and 92.8%) using each of these methods. The PCR method used in this study did not detect all carbapenemase genes; hence, those that tested positive for *bla_KPC_* and *bla_GES_* but negative for other carbapenemase enzymes may process other types of carbapenemase enzyme. Certain bacterial isolates were found to produce carbapenemase enzymes both via the phenotypic method and PCR; other isolates were found to be negative via the phenotypic method and positive via PCR; still other isolates were found to be positive via the phenotypic method and negative via PCR.

However, the absence of gene expression could also be the cause of this. Further explanations for the above could come from substitute resistance mechanisms that mimic the actions of carbapenemase; porin loss, for instance, co-expressing with extended-spectrum β-lactamase and AmpC [30].

Using PCR, the overall prevalence of carbapenemase genes was found to be 92.8% (52/56), with *bla_KPC_* accounting for 80.7% (42/52) and *bla_GES_* for 19.2% (10/52) of the total. The 42 *B. cepacia* isolates that tested positive for carbapenems included 38 *bla_KPC_*(n = 38) and two *bla_GES_* (n = 2); in contrast, four *bla_KPC_*(n = 4) and eight *bla_GES_* (n = 8) were present in the *A. sobria* isolates that tested positive for carbapenems. The present investigation found no positive results for the *bla_IMP_* gene of carbapenem-resistant *A. sobria* or *B. cepacia* isolates. In the present study, *B. cepacia* isolates carbapenems included 40 *bla_KPC_*, in line with Al-Muhanna et al.’s 2020 study which found that the *blaKPC* gene yielded negative results for every *B. cepacia* isolate [4]. Haghighi and Goli, in 2022, found that two *bla_GES_*genes were detected in 83.72% of isolates among gram-negative bacteria like *Pseudomonas aeruginosa* [32]. Shanmugam and associates found that none of *Klebsiella pneumonia* isolates produced *bla_GES_*. The transposable elements that surround the *bla_KPC_* genes, which encode *bla_KPC_*, enable the gene to migrate back and forth between the bacterial chromosome and the transferable plasmid [33]. Difference in resistance to only one of the carbapenems and resistance to both imipenem and meropenem suggest the synthesis of carbapenemase, as this may indicate the existence of another form of resistance mechanism [33,34]. The present investigation found no positive results for the *bla_IMP_* gene of *A. sobria* or *B. cepacia* isolates, consistent with Al-Muhanna et al. who showed that *B. cepacia* isolates gave a negative *bla_IMP_* gene result [4]. According to recent research, mobile gene cassettes inserted into integrons carry two major groups of imported metallo-β−lactamases: VIM and IMP. With the exception of aztreonam, the majority of β-lactam antibiotics are hydrolyzed by metallo-β-lactamases; as a result, most β-lactam antibiotics, including the carbapenems, are ineffective against the many infections that produce these enzymes in high quantities [35,36]. Subsequently, many different gram-negative bacteria isolated from different geographic locations have been shown to carry mobile genetic elements encoding metallo-β-lactamases other than *P. aeruginosa* [37,38]. A number of studies in Asian and European countries, the American countries including Brazil, Canada and the United States, and Australian countries have reported these isolates [39,40].

## Conclusion

To validate the identification of *B. cepacia* complex, the current result showed that *recA* gene molecular identification approach provides a low-cost and highly accurate alternative to the VITEK 2 System. The study suggest that while the isolates exhibited significant rates of resistance to the majority of tested antibiotics, there was a notable predominance of *bla_KPC_*and *bla_GES_* carbapenemase producers among the isolates under investigation. *bla_IMP_* gene was not found in any of the research isolates. The phenotypic approach is a straightforward technique that can identify a wide variety of carbapenemase enzymes; however, it is not specific enough to identify all enzyme types and is unable to identify others. On the other hand, PCR is a more sensitive, quick, and precise technique for identifying particular carbapenemase enzyme types. To find out the ancestry of the isolates, genomic sequencing of the isolates is advised, especially the ones that produce carbapenemase. Future research should focus on using *recA* nucleotide sequence analysis as a quick and repeatable way to detect both newly emerging species and genomovar that are already present in the Burkholderia cepacia complex. To improve our knowledge of the clinical risks connected to each genomovar and novel species within the complex, it will be essential to use this method for the rapid identification of bacterial isolates from patients and other at-risk persons.

## Author Contributions

**Conceptualization:** Mushtak T. S. Al-Ouqaili, Rawaa A. Hussein.

**Formal analysis:** Mushtak T. S. Al-Ouqaili, Rawaa A. Hussein.

**Funding acquisition:** Mushtak T. S. Al-Ouqaili, Rawaa A. Hussein.

**Investigation:** Bushra A. Kanaan, Ahmed T.S. Al-Neda.

**Methodology:** Bushra A. Kanaan, Ahmed T.S. Al-Neda.

**Resources:** Mushtak T. S. Al-Ouqaili, Rawaa A. Hussein.

**Supervision:** Mushtak T. S. Al-Ouqaili.

**Writing – original draft:** Mushtak T. S. Al-Ouqaili, Rawaa A. Hussein.

**Writing – review & editing:** Mushtak T. S. Al-Ouqaili, Rawaa A. Hussein.

## Notes

### Competing Interest Statement

The authors have declared no competing interest.

